# Genetic diversity within diagnostic sputum samples is mirrored in the culture of *Mycobacterium tuberculosis*

**DOI:** 10.1101/2024.01.30.577772

**Authors:** Carla Mariner-Llicer, Galo A. Goig, Manuela Torres-Puente, Sergo Vashakidze, Luis M. Villamayor, Belén Saavedra-Cervera, Edson Mambuque, Iza Khurtsilava, Zaza Avaliani, Alex Rosenthal, Andrei Gabrielian, Marika Shurgaia, Natalia Shubladze, Alberto L. García-Basteiro, Mariana G. López, Iñaki Comas

## Abstract

Culturing *Mycobacterium tuberculosis* (MTB) from tuberculosis cases is the basis for many research and clinical applications. Paradoxically, it is assumed to impose a diversity bottleneck, which, if true, would entail unexplored consequences. The alternative, culture-free sequencing from diagnostic samples, is a promising but challenging approach both to obtain and analyse the MTB genome from the complex sample. This study obtains high-quality genomes of sputum-culture pairs from two different settings after developing a workflow for sequencing from sputum and a tailored bioinformatics pipeline. Our approach reveals that 88% of variants called in culture-free sequencing analysis are false positives due to supplementary alignments, mostly in enriched-sputa samples. Overall, contrary to the bottleneck dogma, we identify a 97% variant agreement within sputum-culture pairs, with a high correlation also in the variants’ frequency (0.98). Our findings extrapolate to all publicly available data, thus demonstrating that in most cases culture accurately mirrors clinical samples.

## INTRODUCTION

*Mycobacterium tuberculosis* (MTB) research from clinical samples usually involves a culturing step to obtain sufficient bacteria for downstream applications, including MTB whole-genome sequencing (WGS) studies. As a consequence, our current knowledge of MTB characteristics, including its biology during infection, evolution, epidemiology, and diagnostics, is largely based on cultured samples ^1,2^. It has been hypothesised that the cultivation procedure may constrain the genetic diversity of MTB, either by selecting for specific variants more suited to in vitro growth as happens for some drug resistance mutations or lineages ^3–5^, or simply due to the bottleneck imposed by the culture inoculum ^6^. If true, this could distort our understanding of bacterial diversity, particularly at the within host level and even affect epidemiological and drug resistance inferences that rely on the presence or absence of a few single nucleotide polymorphisms (SNPs). Direct sequencing from clinical samples is an alternative as it could bypass the potential disadvantages of culturing ^7^. However, the implementation of culture-free sequencing techniques is challenging due to the complexity of the sample matrix with low amounts of mycobacterial DNA and a mix of contaminants including host genetic material ^8,9^.

Previous efforts on culture-free genome sequencing have focused on developing new and affordable protocols for MTB culture-free WGS and assessing their reliability for clinical applications, mainly for AMR diagnosis and transmission inference ^10–15^. Those studies show contradictory results regarding overall genetic diversity comparison between culture-based and culture-free WGS. Some publications have identified no significant differences ^11,12^, while others have reported a reduction in genetic diversity when culturing ^13,16^. These contradictory results probably reflect limitations of the studies, often focused in one single setting and the limited quality of culture-free sequences prevent a proper comparison of genetic diversity between settings. This is specially true to identify low frequency variants which are more likely to suffer from any technical (ie. sample processing) and analytical limitations ^17^. Therefore, the question of whether culture does actually impose such a bottleneck remains largely unsolved despite its importance. The main goal of this work is to determine whether culture reflects the original MTB variability present in the diagnostic samples, typically sputum, making the culture a suitable sample for clinical research.

Here we have put together our own comprehensive dataset including samples from two settings differing in tuberculosis (TB) incidence, burden of AMR, and HIV co-infection; and achieving sufficient sequencing depth to properly compare genetic diversity, especially regarding low-frequency variants. We successfully sequence 61 high-quality sputum-culture pairs from Georgia and Mozambique. For the sputa, we implemented a culture-free WGS approach based either in direct (dWGS) or bait-enrichment (eWGS) sequencing, depending on the amount of MTB DNA. In addition we carry out experimental benchmarking to detect major sources of artifactual genetic variation in culture-free approaches. We also reanalyze available datasets ^10,13,16^ to generalise our results across settings and sequencing approaches. Importantly, we develop a tailored analysis workflow to address the absence of standardised laboratory protocols and bioinformatic pipelines, carefully considering artefacts beyond the role of potential contaminants in the sample. Our customised workflow is capable of demonstrating that culture accurately reflects sputum diversity in all the evaluated settings, albeit with some individual exceptions. Our results indicate that the current knowledge of *M. tuberculosis* diversity based on culturing methods is robust and therefore the genetic diversity observed in culture generally mirrors that present in the diagnostic sample.

## RESULTS

### Selection of sputum-culture pairs

Out of the initial 95 sputum-culture pairs available, 80/95 (84.2%) were suitable for WGS since 4/95 (4.2%) cultures did not grow and 11/95 (11.6%) sputa were MTB negative according to qPCR results (Cq>35). Given their quality, in terms of Cq and %MTB DNA, 48/80 (60%) sputa underwent dWGS and 32/80 (40%) were enriched before sequencing (**Fig. 1a**; Methods **Fig. 5**). Percentage of MTB was obtained before and after the enrichment step. We observed that sputa with an initial MTB% within 0.5-10% reached 25.5-98.4% after the enrichment, while those containing initially 10-25% of MTB reached more than 90% of MTB (Methods **Fig. 6**).

**Fig. 1:**
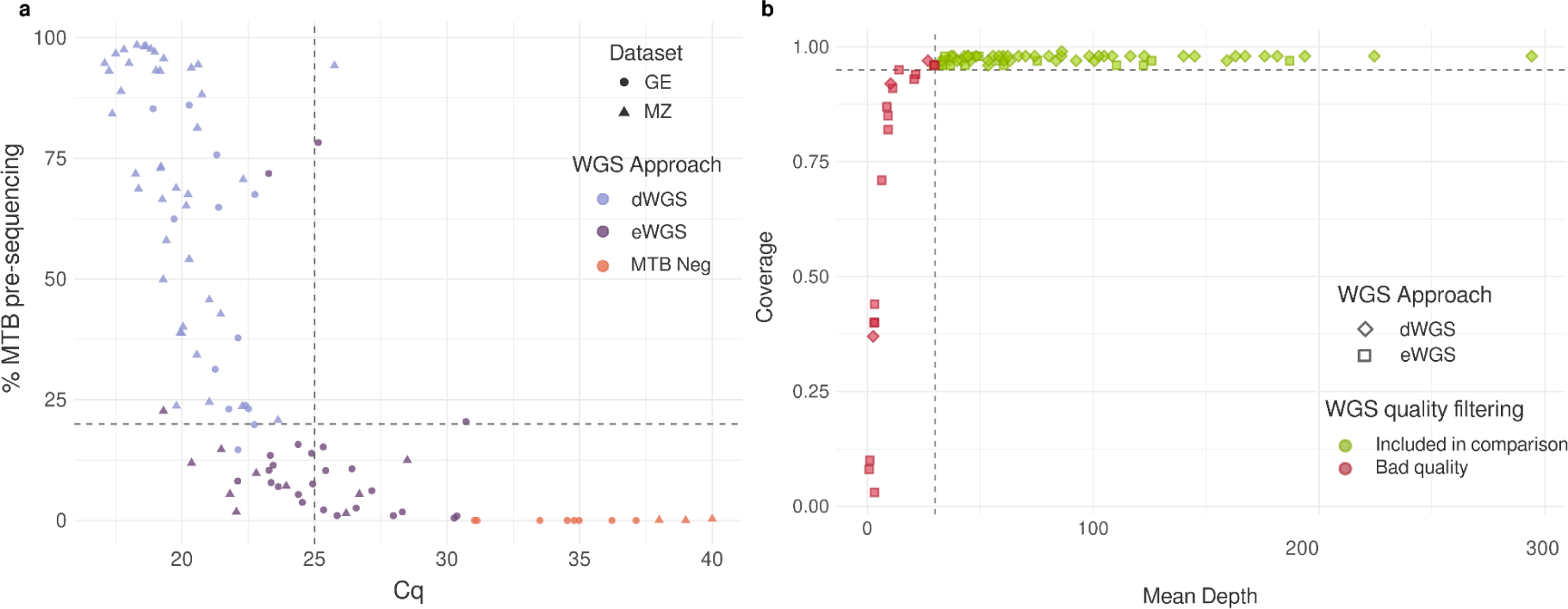
Evaluation of sputum samples for sequencing. **a,** qPCR Cq vs %MTB. 95 sputa are each one represented by a point. %MTB was obtained by performing a pre-sequencing run in order to determine if each sputum contained enough MTB DNA to be sequenced directly (light purple) or if it required a previous enrichment step (dark purple). Sputum samples in orange were considered negative (Cq>35 and %MTB <1%). Shape indicates sample origin: triangles for Mozambique, dots for Georgia. Dashed lines represent thresholds to decide which WGS approach to follow, the horizontal line highlights MTB% = 20%, the vertical line highlights Cq = 25. **b,** Mean depth vs coverage. 86 sputa sequenced with enrichment (eWGS, square) or directly (dWGS, diamond). Colour represents the sequencing quality, good quality samples (the ones used for comparison analysis) are in green and bad quality samples in red. The dashed lines represent the coverage and depth cut-off values to consider a good quality sample, the horizontal line highlights a coverage = 0.95 and the vertical line highlights depth = 30X.

A total of 19 sputa sequences did not meet the minimum quality criteria (30X depth and 95% coverage) (**Fig. 1b**), thus these pairs were excluded from further analysis. The sequencing performance of the remaining 61 pairs was: median depth 48X for eWGS (30-157X), 74X (33-302X) for dWGS and 114X (46-347X) for cultures. Median genome coverage, at a minimum of 20X depth, was above 96% in all sputa and cultures (**Table 1**). The samples’ sequencing workflow is detailed in the Methods section **Fig. 5**. All samples’ information is available at **Table S1.**

**Table 1:** Summary of sequencing results of the 61 paired samples.

|  | Total pairs = 61 |  |  |
| --- | --- | --- | --- |
|  | eWGS | dWGS | Culture WGS |
| **Total WGS** | 16 | 45 | 61 |
| **Median Depth** | 48.00 | 74.00 | 114.00 |
| **Q1** | 33.75 | 47.00 | 94.00 |
| **Q3** | 76.25 | 120.00 | 127.00 |
| **IQR** | 42.50 | 73.00 | 33.00 |
| **min depth** | 30.00 | 33.00 | 49.00 |
| **max depth** | 157.00 | 302.00 | 347.00 |
| **Median Coverage** | 0.96 | 0.98 | 0.98 |
| **Q1** | 0.96 | 0.97 | 0.98 |
| **Q3** | 0.97 | 0.98 | 0.98 |
| **IQR** | 0.01 | 0.01 | 0.01 |
| **min coverage** | 0.96 | 0.96 | 0.97 |
| **max coverage** | 0.98 | 0.99 | 0.98 |

**Abbreviations:** Q - quartile, IQR - Interquartile range, min - minimum, max - maximum

### Tailored SNP calling pipeline for culture free sequencing analysis

To determine if culture-free sequencing approaches introduced a bias during the variant calling, we compared fixed and low-frequency variants detected in trios of eWGS, dWGS and culture-WGS from the same TB case in three suitable examples. After variant calling, we implemented a “recovery” step in which all variants not passing filters in one sequencing approach, but accurately called in another approach, were re-included or “rescued” in the analysis. Our rationale was that culture-free WGS is performed in samples with low amounts of MTB DNA, hence increasing the chance of variants not passing the filters due to lower sequencing depths. Before applying our customised calling filters, we identified up to 22 variants only detected in eWGS, all of them at low frequencies (median=22%, range=10-54%). Exclusive variants were also detected in dWGS (up to 8) and in culture-WGS (up to 3) but the magnitude of the differences were minor compared to eWGS (**Fig. 2a** Venn diagrams in blue). We screened manually the discrepant variants in the eWGS approach, and identified that all of them appeared in supplementary alignments (**Supplementary material Fig. S2**). A supplementary alignment is a read segment split from the primary read and aligned to a different region of the genome that can produce false positive calls in the variant calling step ^18,19^. After discarding supplementary alignments, most discrepant SNPs disappeared giving a concordance of 99-100% in the variant calling between the three sequencing approaches (**Fig. 2a**).

**Fig. 2:**
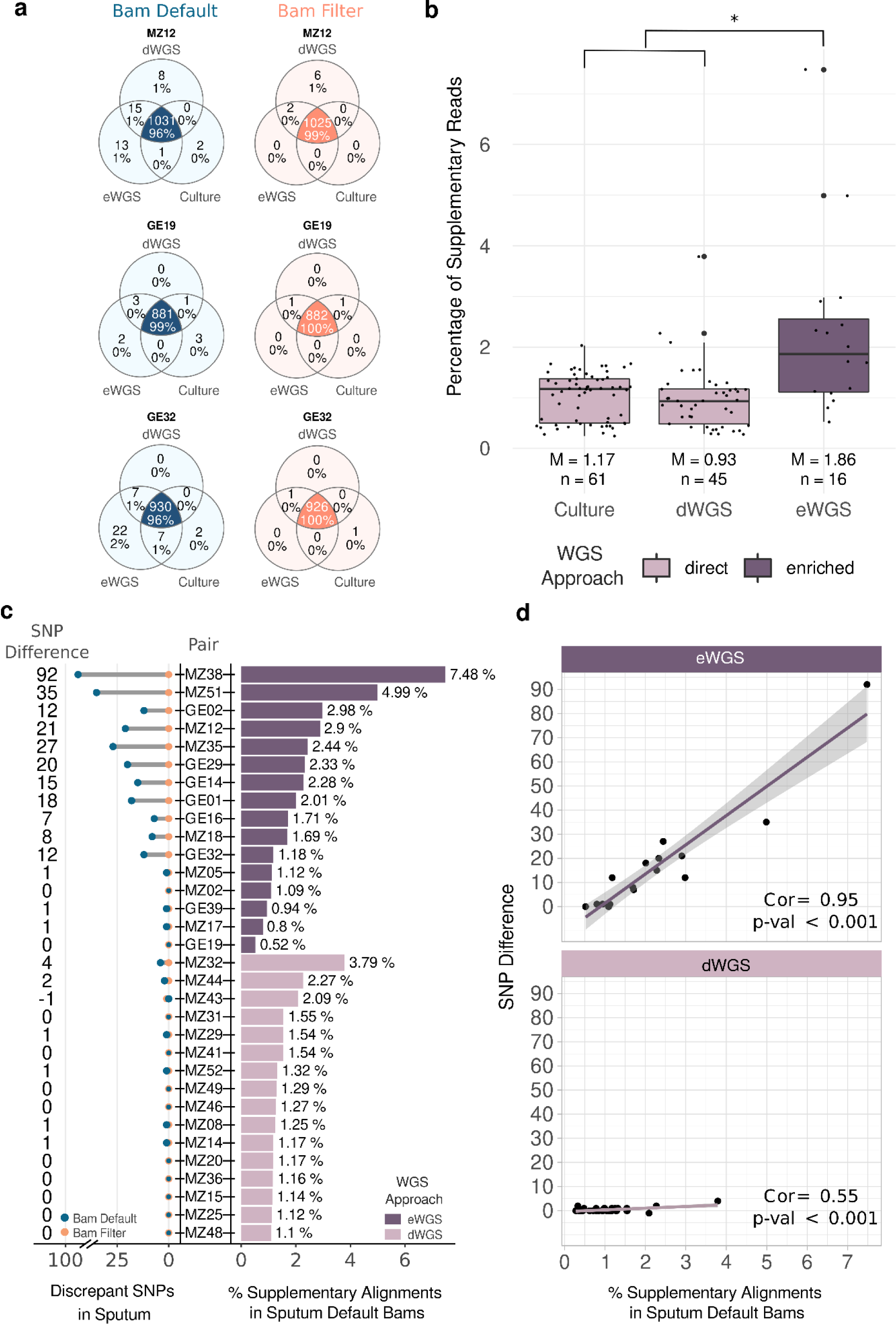
Analysis of supplementary alignments. **a**, Venn diagrams of the comparisons between trios of direct sputum (dWGS) - enriched sputum (eWGS) - culture WGS. Amount of exclusive and common variants, together with the percentages, are denoted. Blue Venn diagrams represent comparisons of variant calls from default unfiltered bams (including supplementary alignments. Orange Venn diagrams represent comparisons of variant calls from filtered bams. **b,** Comparison of the amount of supplementary alignments between direct sequencing (sputum dWGS and culture WGS, light purple) and enriched sputum samples (eWGS, dark purple). Median (M) and the total amount of samples (n) are shown. Asterisk (*) highlights a significant p-value (<0.001). **c,** On the left there is the comparison of the amount of discrepant SNPs exclusive in sputum, either dWGS and eWGS, before and after supplementary alignments filtering. Dots colours stand for variant calls from bam files before discarding supplementary alignments (in blue) and after filtering them (in orange). The x axis is discontinued. The right part shows the percentage of supplementary alignments in sputum files, either dWGS and eWGS. Colour represents the sequencing approach for sputum samples, dWGS sputa appear in light purple and eWGS in dark purple. Plot c contains 32/61 pairs, the 16 ones containing a higher percentage of supplementary alignments in each eWGS and dWGS. Samples are ordered from the highest to the lowest amount of supplementary alignments. The complete version containing the 61 pairs can be seen in **Supplementary Material Fig. S3. d,** Correlation between the percentage of supplementary alignments and the amount of SNPs removed when discarding supplementary alignments form sputum bam files (represented as SNP difference and calculated as follows: discrepant SNPs exclusive in sputum in Default Bams - Filtered Bams). Pearson correlation coefficients and p-values are shown.

The extended analysis to all the 61 sputum-culture pairs demonstrated that overall the 88.5% (307/347) of the discrepant variants between sputa (either eWGS or dWGS) and paired cultures were false positives introduced by supplementary alignments. Particularly, eWGS sputa accumulated significantly higher false positive variants than dWGS sputum and cultures (Wilcox test, p-value < 0.01. **Fig. 2b**) due to a higher amount of chimeric reads causing supplementary alignments. After discarding them from the bam files, we identified a mean of 18.3 false positive SNPs (range 0-106 SNPs corresponding to 1-7.4%) per eWGS-culture pair (**Fig. 2c, Supplementary material Fig. S3**) whereas, dWGS-culture pairs showed a lower mean value of 0.3 (1-5 SNP corresponding to 0.3-5%) false positive (**Fig. 2c, Supplementary material Fig. S3**). Notably, a high, positive and significant correlation (corr 0.95, p-value < 0.001) was obtained between supplementary alignments and false positive SNPs for eWGS, whereas, in the case of dWGS, the correlation was lower and with limited impact on variation (**Fig. 2d**). Supplementary alignments accounted for only 1.2% of the reads on average (range: 0.2-7.5; dWGS: 1.0%, eWGS: 2.3%), therefore not impacting the depth and genome coverage (**Supplementary Table S1**).

### Sputum-culture genetic diversity comparison

After applying our tailored SNP calling pipeline we evaluated the genetic diversity between sputum and culture. Overall, a mean of 913 (range 770-1063), representing 97% of the total SNPs detected, were common in sputum and culture. A small proportion of rescued SNPs was observed with a mean of 24 (range 5-69) representing less than 3% of total SNPs (**Table 2**; **Fig. 3a; Supplementary material Fig. S4**). Regarding exclusive variants, we also observed a small proportion of sputum-exclusive SNPs (mean 1, range 0-10) and culture-exclusive (mean 1, range 0-11). In other words, most of the sputum-culture pairs (49/61, 80.3%) did not display any discrepancies in sputum; 9/61 (14.8%) presented 1-5; and only 3/61 (4.9%) had between 5 and 10 sputum-exclusive SNPs (**Fig. 3b**).

**Fig. 3:**
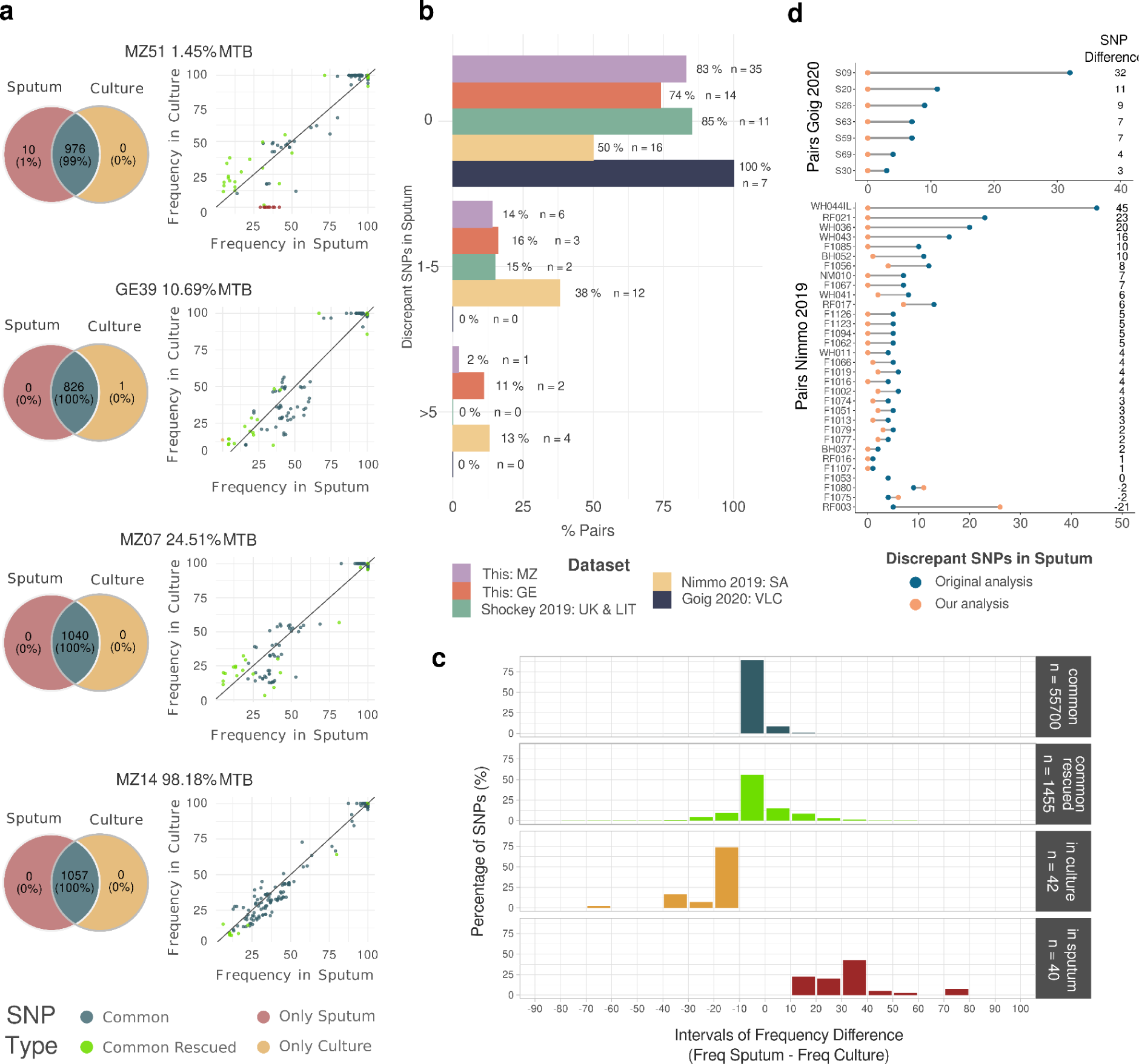
Comparison of variants between sputum-culture pairs. **a,** Comparison of the amount of common and exclusive variants (Venn diagrams on the left) and comparison of frequency of variants in sputum (dWGS or eWGS) and culture (on the right). Colours represent common or exclusive variants. Percentage of MTB reads in the dWGS is shown above each plot. The complete figures containing the 61 pairs can be seen in **Supplementary Material Fig. S4 and Fig. S5**. **b,** Analysis of sputum-exclusive variants in this dataset and published ones. Percentage of pairs in each dataset with 0, 1-5 or more than 5 sputum-exclusive SNPs (all were not fixed variants). Colours represent the dataset. Abbreviations of countries/regions stand for: MZ-Mozambique, GE-Georgia, UK-United Kingdom, LIT-Lituania, SA-South Africa, VLC-Valencia (Spain). **c,** Histogram of the difference of frequency between variants obtained in sputum versus culture (frequency in sputum - frequency in culture). Colours represent whether the variants are common or exclusive. Total number of variants is shown in grey boxes (n). **d,** Differences of sputum-exclusive SNPs published in the original paper (in blue) versus the ones found by running our pipeline (in orange) for Goig *et al.* 2020 ^10^ (upper panel) and Nimmo *et al.* 2019 ^16^ (bottom panel) datasets.

**Table 2:**
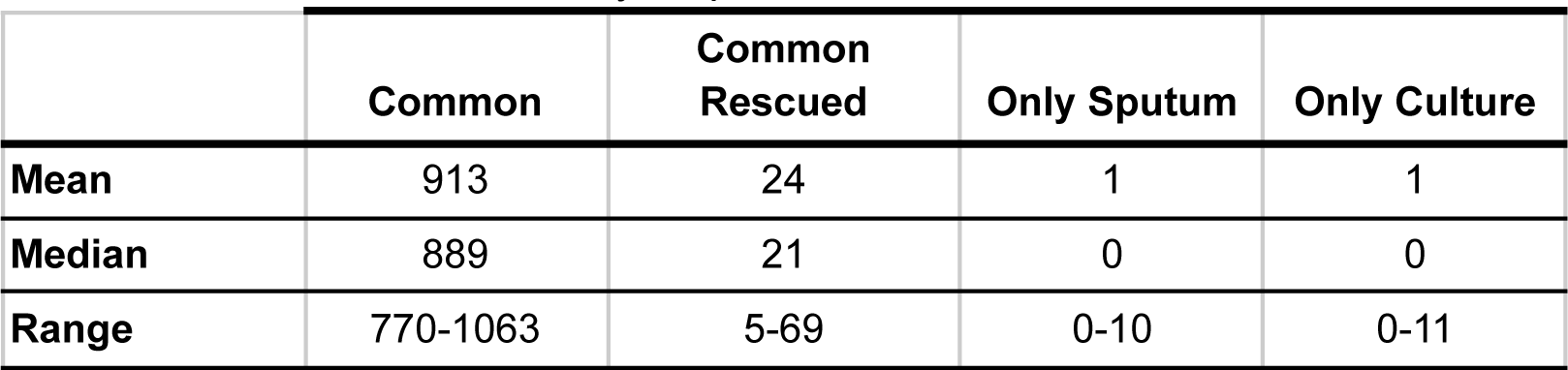
Counts of common and discrepant variants in 61 pairs. Exclusive SNPs shown were obtained after the recovery step.

|  | Common | Common Rescued | Only Sputum | Only Culture |
| --- | --- | --- | --- | --- |
| Mean | 913 | 24 | 1 | 1 |
| Median | 889 | 21 | 0 | 0 |
| Range | 770-1063 | 5-69 | 0-10 | 0-11 |

It could be argued that culture-free WGS will be more relevant for determining the frequency of SNPs rather than just their presence/absence. The correlation between the frequencies of SNPs observed in both sputum, either dWGS and eWGS, and culture (common SNPs) was 0.98 (0.93-0.99, p-value<0.001) (**Fig. 3a; Supplementary material Fig. S5**). By going deeper, we observed that most of the common SNPs (98%) had a small frequency difference within both paired samples (ranging 0-10%) with higher frequencies in culture than in sputum (**Fig. 3c**). As expected, these differences were slightly higher for rescued SNPs with 70% of them ranging 0-10% frequency and a 28% ranging 11-30% frequency (**Fig. 3c**).

Regarding the exclusive variants, we observed a total of 40 sputum-exclusive SNPs distributed among 12 sputa, with 17 out of 40 (42%) falling within a frequency interval of 30-40%. For those culture-exclusives, which were present in 12 cultures, 73.8% (31/42) were found within a frequency range of 10-20% (**Fig. 3c**).

The results for discrepant variants suggest that, in some cases, variants with intermediate frequencies (between 30-80%) in sputum can be missed in culture. In any case, the number of exclusive variants present only in sputum or in cultures were similar and low compared to those commonly called. To highlight, no fixed-SNP (>90% frequency) was detected within them.

### Generalisation of results

Finally, we extended our analysis to publicly available sputum-culture sequencing data from three different clinical settings ^10,13,16^. In those, authors described differences in genetic diversity between sputum and culture pairs. We only analysed those sequences meeting the quality criteria (see **Methods**) for comparisons. After applying our pipeline, we observed that 88% of pairs differed in less than 5 SNPs (**Fig. 3b**), all of them below 90% frequency. These results are very similar to the ones obtained for the dataset generated in this study (samples from Georgia and Mozambique; **Fig. 3b**). For those studies with the number of discrepant SNPs available in the publication (Goig *et al.* 2020 ^10^ and Nimmo *et al.* 2019 ^16^), our pipeline reduces the number of discrepant SNPs from an average of 10.4 SNPs (median 7, range 4-32) to 0 SNPs in all 7/7 sputum-culture pairs of Goig *et al.* ^10^ dataset, and from an average of 8.7 SNPs (median 5, range 1-45) to 1 SNPs (median: 0, range 0-7) in 28/32 pairs from Nimmo *et al.* ^16^ dataset (**Fig. 3d**). Contrary to the Goig *et al*.^10^ dataset, in three samples of Nimmo *et al*.^16^ dataset, the number of SNPs did not decrease before and after applying our tailored pipeline, and there was even an increase in one of the pairs.

Overall, the results from our dataset suggest that almost all the genetic variability present in the original sputum is represented by culture both in terms of presence/absence of SNPs but also in terms of SNP frequency correlation within the samples. Re-analysis of available datasets also points to a high sputum-culture concordance after discarding false-positive variants, mostly due to the bioinformatic pipeline implemented.

### Phylogenetic classification and drug-resistance profile

We obtained a 100% concordance between sputum and culture lineage (L) prediction. We identified 31/61 L4 strains (20 from Mozambique and 11 from Georgia) and 16/61 L2 (8 strains from each dataset). The remaining strains from Mozambique belonged to L1 (13/61 samples) and L3 (1/61 sample). Concordance at lower taxonomic levels according to Coll *et al.* classification ^20^ was also 100%. Sublineage classification is shown at **Supplementary Table S2**.

All the 61 sputum samples matched with their paired cultures when constructing a Maximum likelihood tree (**Supplementary material Fig. S1**) obtaining a 0 SNP in all pairs (fixed-SNP >90% frequency) pairwise distance. Two samples from Mozambique were in the same transmission cluster (measured at a 5 SNP distance cut-off) which was equally detected when culture and sputum genomes were analysed separately.

Regarding AMR-conferring variants, the agreement between the resistance profile predicted in sputum and culture was 100%. We obtained 10/61 pairs with at least one AMR-conferring mutation (6 from Georgia, 4 from Mozambique): 7/10 mono or poly-resistant, 2 MDR and 1 pre-XDR. All these were fixed-SNPs in both sputum and culture. No low-frequency AMR-conferring variants were found. See **Table S3** for resistance SNPs information.

## DISCUSSION

In this study we explore the role of culturing from sputum diagnostic samples as a bottleneck for MTB genetic diversity. In terms of SNPs’ presence/absence, both fixed and minority variants, after comparing the genomic variability, most culture-sputum pairs (91.8%, 56/61) showed a difference of less than 5 (0-3) SNPs. Strikingly, the correlation of the SNPs’ frequencies was very high (0.98, p-value < 0.001). Our findings show that culture mirrors the genetic diversity presence in the sputum, considering both presence/absence of SNPs and their frequencies. We corroborate our results by reanalyzing previously published datasets, suggesting that any apparent contradictions were likely due to differences in bioinformatic pipelines. Importantly, our findings were consistent across datasets, encompassing samples from different clinical settings and laboratories.

One key finding of our study was the detection of false-positive variation driven by chimeric reads most likely produced during sample enrichment and library preparation steps. While this is a common issue, we observed that they were significantly higher in the enriched sputa as compared to direct sputa sequenced. The presence of chimeric reads was potentially due to the low concentration of DNA in sputa. Considering that not all DNA was from MTB, the low quantity of target DNA could lead to non-specific amplifications in both library preparation and PCR reactions carried out during the enrichment. Such amplifications likely generate chimeric reads, resulting in supplementary alignments during the mapping and, therefore, false variant calls. Furthermore, a high correlation between supplementary alignments and false-positive SNPs was obtained for eWGS sputa. Removing supplementary alignments reduces by more than 80% the discrepant variants between sputum and culture without impacting depth and coverage.

Another source of discrepancies is driven by differences in depth throughout the genome, which is more relevant in culture-free WGS approaches. To solve this, here we implemented a recovery step to rescue the false negative variants present in the sputum. It is worth noting that the absence of culture could have resulted in the oversight of these variants in the sputum. However, it was observed that the rescued SNPs only make up less than 3% of the total SNPs in the pair. Overall, these findings unequivocally indicate that achieving an accurate representation of the original genetic diversity depends on sequencing depth and appropriate bioinformatic filters.

The main aim of this study was to assess whether culturing distorts the genetic diversity of the bacterial population present in sputum. By performing an enrichment step, we succeed in increasing the MTB DNA by 26.5-fold on average, allowing high-quality sequences of sputa with as low as 1% of MTB DNA, similar to the values obtained by Mann *et al.* ^21^. Since we have not explored the potential of culture-free WGS as a diagnostic tool for TB, we performed as many runs as necessary to achieve sufficient coverage and depth to reach our goal, regardless of the sequencing cost per sample. With this effort we were able to perform culture-free sequencing, with high quality, in 76% (61/80) of diagnostic samples, suggesting that further improvements of the technique will be needed as shown by others ^15^. In fact, while previous studies showed that smear-negative and scanty sputa could be enriched and sequenced occasionally ^10–13^, a formal testing of the limit of detection is still needed. Meanwhile, intermediate alternatives like targeted next-generation sequencing tools are finding a room in the TB diagnostics pipelines ^22^.

Understanding that the diversity of *M. tuberculosis* is reflected in both sputum and culture has an impact in TB research, since the overwhelming majority of studies today rely on culture sequencing to assess diversity, even in those cases where surgery ^23^ or post-mortem ^24^ samples are interrogated. Our analysis suggests that diversity is well represented in those studies and that substantial variation is not missed when culturing. However, as a limitation, in this study we have not been able to analyse the detection of genotypes carrying AMR mutations (or any other mutation) with a fitness cost in culture. Most of our samples are drug susceptible and our analysis has not identified any heteroresistance. Thus, the high correlation observed in our sputum-culture pairs may not apply in situations where a subpopulation has a mutation with associated fitness cost. Mutations such as these are known to cause growth delay and affect culture-based diagnostics, particularly in the case of rifampicin-resistance conferring mutations ^4,25^. Similarly, this limitation also applies to cases where lineages with different growth rates coexist, potentially affecting the identification of polyclonal infections, as it has been previously suggested for Lineage 6 and animal strains ^5^. Identification of those scenarios requires specific analysis ^23^ out of the scope of this work. For some cases, particularly in the datasets from Georgia and South Africa ^16^, we identified a higher disagreement between sputum and culture. It likely be due to the presence of unknown fitness-cost associated mutations or to polyclonal infections with genetically close strains, which are common in both settings ^26^. Nevertheless, our dataset reflects the reality of many TB cases, which are from single infections and carry drug resistance mutations that do not cause growth problems in culture.

In conclusion, our results highlight the importance of evaluating and applying appropriate filtering steps when sequencing complex samples, such as sputum, in order to detect and discard sources of false variation. After developing and applying a tailored bioinformatics pipeline, we show that culture accurately captures the genetic diversity present in diagnostic samples and this is true across settings and laboratories. On one hand, from a diagnostic point of view our results reflect the long road ahead towards whole genome-based diagnostics, as highlighted in recent publications ^15^. On the other hand, from a research point of view our results support the large body of work based on culture sequencing.

## ONLINE METHODS

### Dataset

We received sputum-culture paired samples from the Centro de Investigação em Saúde de Manhiça (Mozambique) and the National centre for Tuberculosis and Lung Diseases located in Tbilisi (Georgia). To ensure that the bacillary load present in sputum samples was enough for sequencing, we selected 50 sputum samples from Mozambique graded High/Medium according XpertUltra result and 3+/2+ according to smear microscopy result. Regarding Georgia, we processed all 45 sputum samples received due to the lack of bacillary load information. In summary, we processed 95 sputum-culture paired samples.

Sputum samples were homogenised and decontaminated in origin countries. In Mozambique, they performed the N-acetyl-L-Cysteine-sodium hydroxide (NALC-NaOH) or Kubica method ^27^, while in Georgia they followed the modified Petroff protocol, that uses 4% NaOH, which was validated under supervision of the WHO Supranational TB Reference Laboratory in Antwerp ^28^.

The paired cultures were grown in 7H11 solid media, supplemented with OADC and glycerol to ensure a high amount of bacteria for sequencing.

All samples, cultures and sputa, were received and processed at FISABIO biosafety level 3 (BSL-3) facilities in Valencia, Spain.

### Samples’ Processing

#### DNA extraction from sputum samples

DNA extraction from sputum leftovers was based on a differential cell lysis procedure to remove non-MTB contaminant DNA using MolYsis basic5 kit (Molzym, Germany). We followed a modified version of the original protocol that entailed an initial lysis of non-mycobacterial cells to remove contaminant DNA, followed by an inactivation of MTB cells at 95°C for 15 minutes and a mechanical cell disruption using FastPrep. DNA precipitation steps were performed using ethanol, sodium acetate and Glycoblue (Methods **Fig. 4**).

**Fig. 4:**
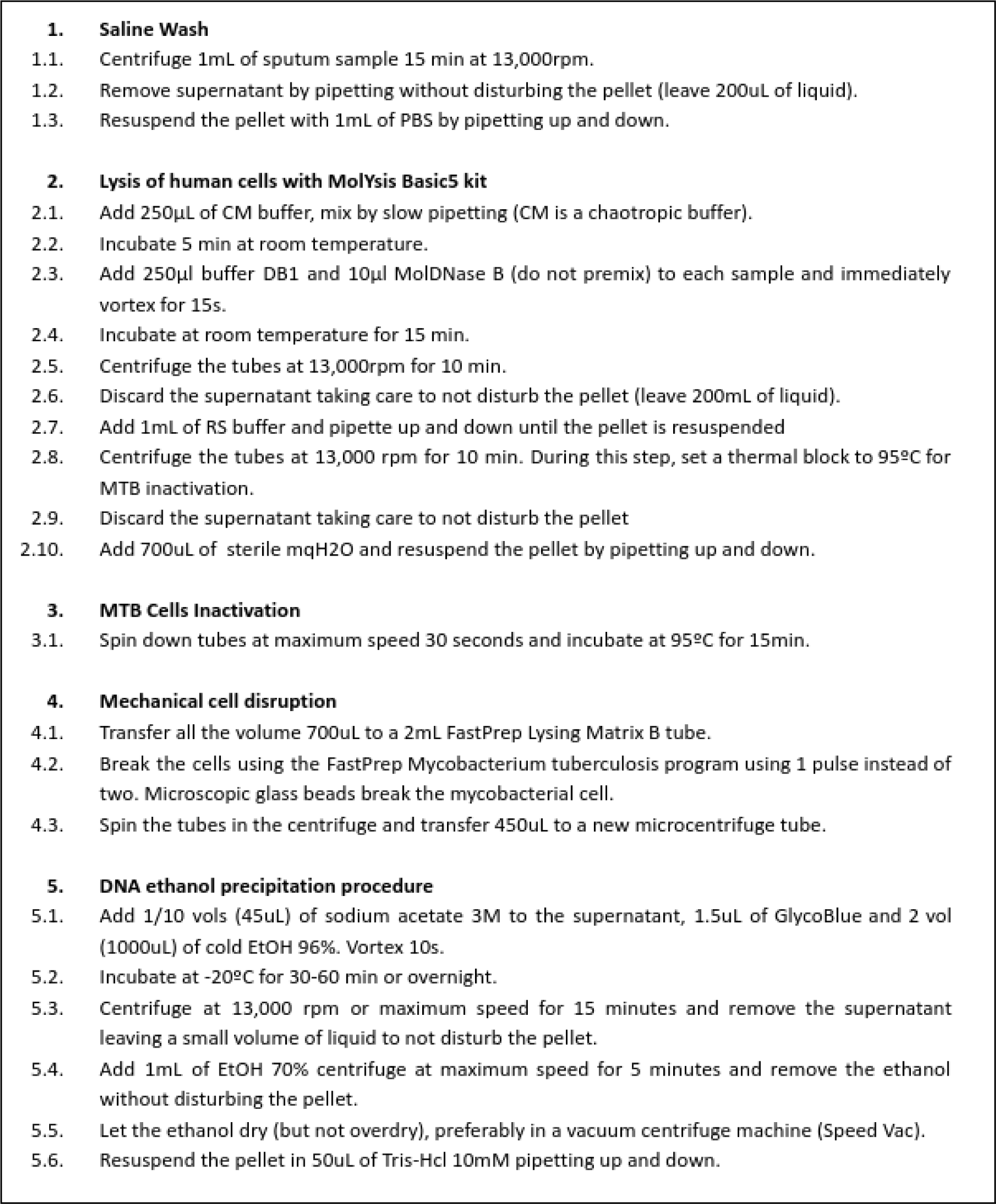
Detailed protocol for DNA extraction protocol from sputum samples (Molysis5 basic adapted protocol).

#### DNA extraction from culture samples

Cultures were heat inactivated at 90°C during 30 minutes and then MTB DNA was extracted following the standardised CTAB protocol ^29^ based on an overnight cell wall digestion with lysozyme, followed by incubation steps with proteinaseK, SDS, CTAB and NaCl and a final DNA precipitation step with isopropanol. All bacterial cultures and DNA extraction steps were performed in a BSL-3 laboratory.

#### qPCR Conditions

We assess the concentration of MTB in sputum samples to decide the sequencing approach. The qPCR was performed in a total volume of 20 uL including 10 uL of Kapa Fast Probe Master Mix 2X, 2 uL of forward and reverse primers mix 2.5 uM, 0.6 uL of probe 10 uM; and 1 ng of DNA. We used DNA normalised to 0.5ng/uL. The qPCR assay consisted on the amplification of a 65 bp region within the *Rv2341* gene using the following primers: Forward-GCCGCTCATGCTCCTTGGAT, Reverse-AGGTCGGTTCGCTGGTCTTG, Probe-TGAGTGCCTGCGGCCGCAGCGC ^30^.

#### Sequencing selection

We prepared NexteraXT (Illumina) libraries for all samples. Libraries from cultures were sequenced directly. Regarding sputum samples, we classified libraries for dWGS (direct) or eWGS (enriched) depending on Cq and %TB obtained in pre-sequencing runs as described in ^10^. Therefore, we performed a qPCR targeting *Rv2341* gene (previously described ^10,30^) to quantify the amount of MTB DNA. Sputa that obtained a Cq value above 35 (equivalent to less than 2 genome copies) were considered negative (not suitable for sequencing) (See ***qPCR conditions*** section). Libraries were prepared for all MTB positive samples (Cq<35) and a pre-sequencing run was performed to estimate the percentage of MTB. According to qPCR and pre-sequencing %MTB result we performed dWGS in sputa obtaining a Cq<25 and more than 20% of MTB and the rest sputum samples underwent an enrichment step before sequencing. The enrichment step consisted of a MTB DNA capture using RNA biotinylated baits (See ***Enrichment step*** section). See the flowchart in Methods **Fig. 5**.

**Fig. 5:**
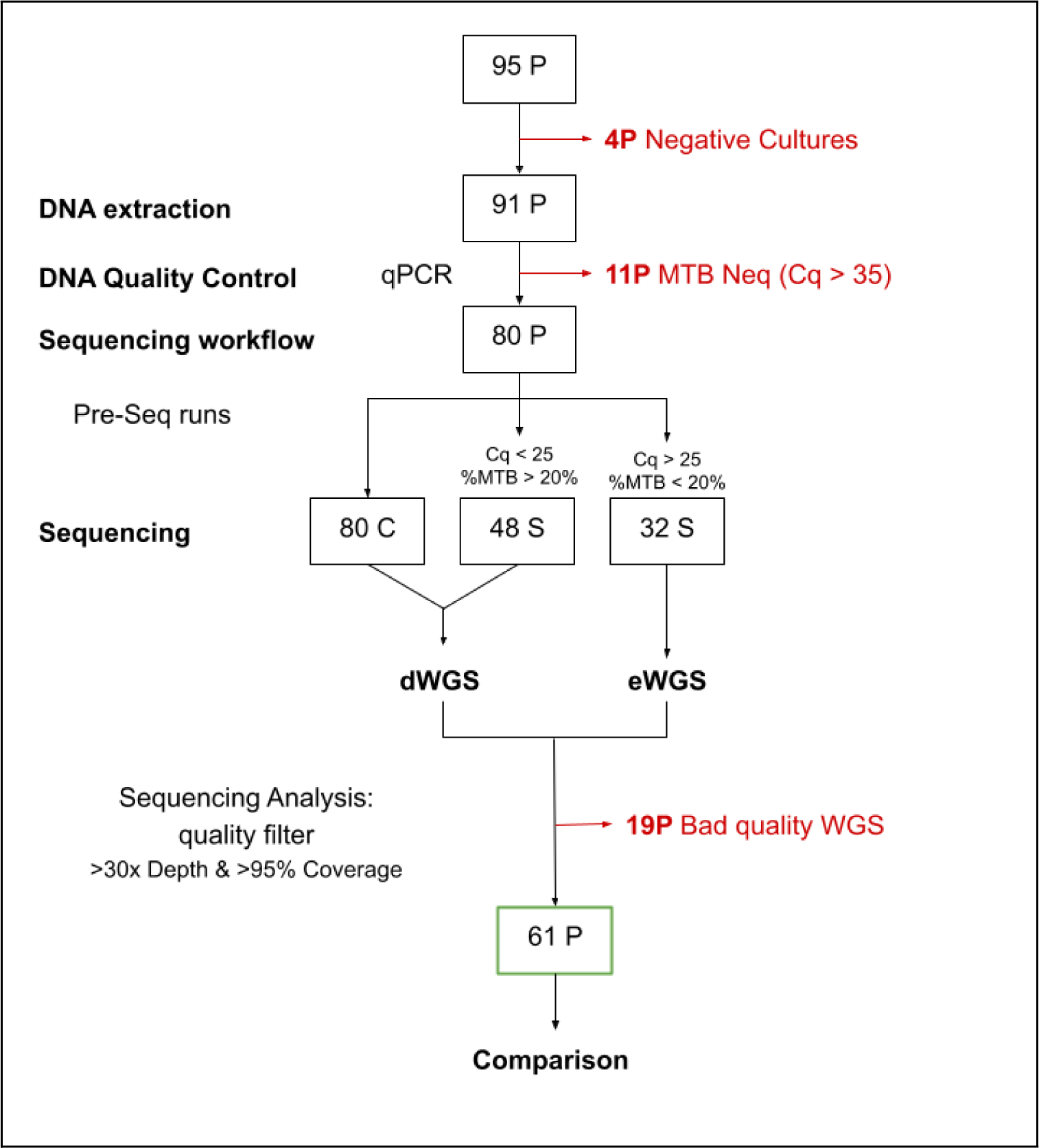
Sequencing workflow. Abbreviations: C-Culture, S-Sputum, P-Pair, dWGS-sputum samples not enriched, eWGS-sputum samples enriched.

WGS was performed in Illumina Miseq (2×300bp) or NextSeq (2×150bp) platforms. Samples were sequenced until a minimum of 30X depth was reached. Sequences are available at European Nucleotide Archive (ENA) under project accession number: PRJEB64897. Detailed information about the samples can be found at **Supplementary Table S1**.

#### Enrichment Step

We used myBaits kit (Arbor Biosciences) to perform the hybridization MTB DNA capture step following the myBaits protocol Version 4.01. This protocol is based on a hybridization of already prepared whole genome sequencing libraries (prepared using NexteraXT kit) with RNA biotinylated baits at 65°C overnight (at least 24h). During this step, baits hybridised to denatured MTB DNA. Then, most non-captured DNA is discarded by different cleaning steps with streptavidin-coated magnetic beads. Finally, the MTB library is amplified by a post-capture PCR in a 50uL reaction containing 25uL of KAPA HiFi HotStart ReadyMix PCR Kit (Roche), 200nM of Illumina sequencing primers P5 and P7; and 15uL of captured library. Post-capture PRC conditions are the following: 15 cycles of 20 seconds to 98°C, 30 seconds to 65°C and 1 minute to 72°C.

The RNA-biotinylated baits panel was designed and developed by Arbor Biosciences using as reference the inferred ancestor genome of the MTB (NC_000962.3), genetically equidistant to all the MTB lineages.

We evaluated the enrichment step by obtaining the fold-change of the percentage of MTB reads between the 32 sputum samples before and after the enrichment. The median fold-change was 10.96 (Methods **Fig. 6**). In other words, sputa containing an initial MTB% within 0.5-10% got a 64.8% (average) while those with an initial MTB% of 10-25% reached 94.3% after undergoing the enrichment step (Methods **Fig. 6**).

**Fig. 6:**
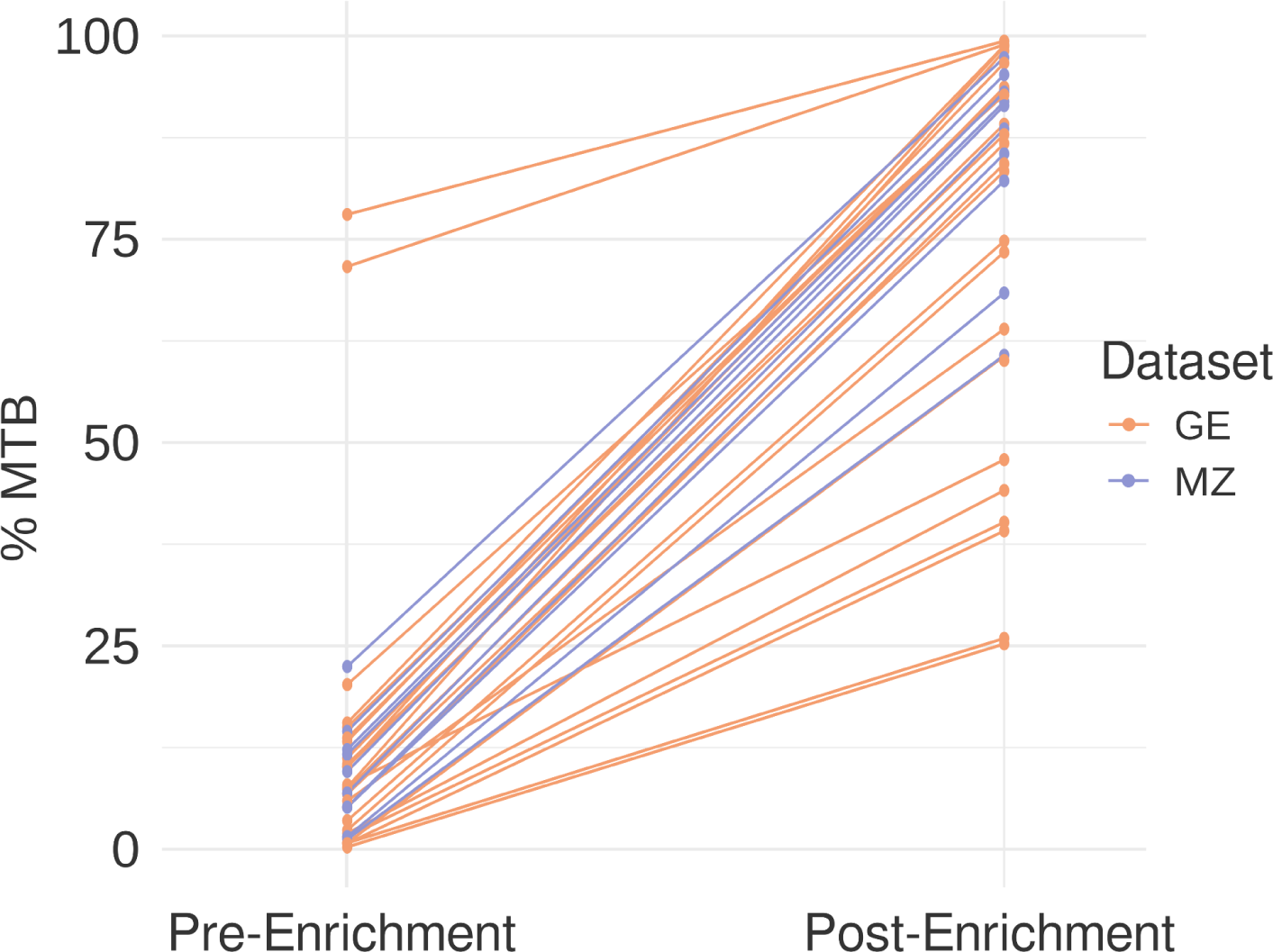
**Comparison of percentage of MTB** before and after the enrichment step. Colours represent samples’ origin.

### Bioinformatics Analysis

#### Core analysis

Analysis was performed using our routine pipeline available at https://gitlab.com/tbgenomicsunit/ThePipeline for both culture and sputum samples. First, reads were trimmed by quality with FastP ^31^; non-MTB reads were discarded by using Kraken ^8,32^. MTB reads were mapped to the reference ancestor genome (NC_000962.3) ^33^, which is genetically equidistant to all lineages, using BWA (version 0.7.10-r789) ^18^.

Samples with a median depth below 30X and less than 95% of the genome covered were discarded for downstream variant comparison analysis. For the remaining good quality samples, variants were called with VarScan (version 2.3.7) ^34^ and Samtools v1.15 ^35^ by applying stricter coverage and frequency filters for culture samples than for sputum samples (variants called in at least 3 reads, in both strands for sputum; variants appearing in 6 reads, in at 10X depth and in both strands, for culture) (parameters for VarScan are shown in Methods **Fig. 7**). For the genetic diversity comparison analysis, we discarded SNPs appearing in high density regions with GATK (version 4.0.2.1) ^36^ or genomic regions that are known to be challenging for short-read mapping such as repetitive genes or mobile elements (PE/PPE gene families) ^37^ by using a customised Python2 (version 2.7.5) script. Highly conserved genes such as *rrs* and *rrl* were also discarded for the comparisons to avoid false positive variability coming from non-MTB reads not discarded by Kraken. We also obtained Pearson’s correlation of the variant frequency obtained in each sputum-culture pair (the mean, median and the range).

**Fig. 7:**
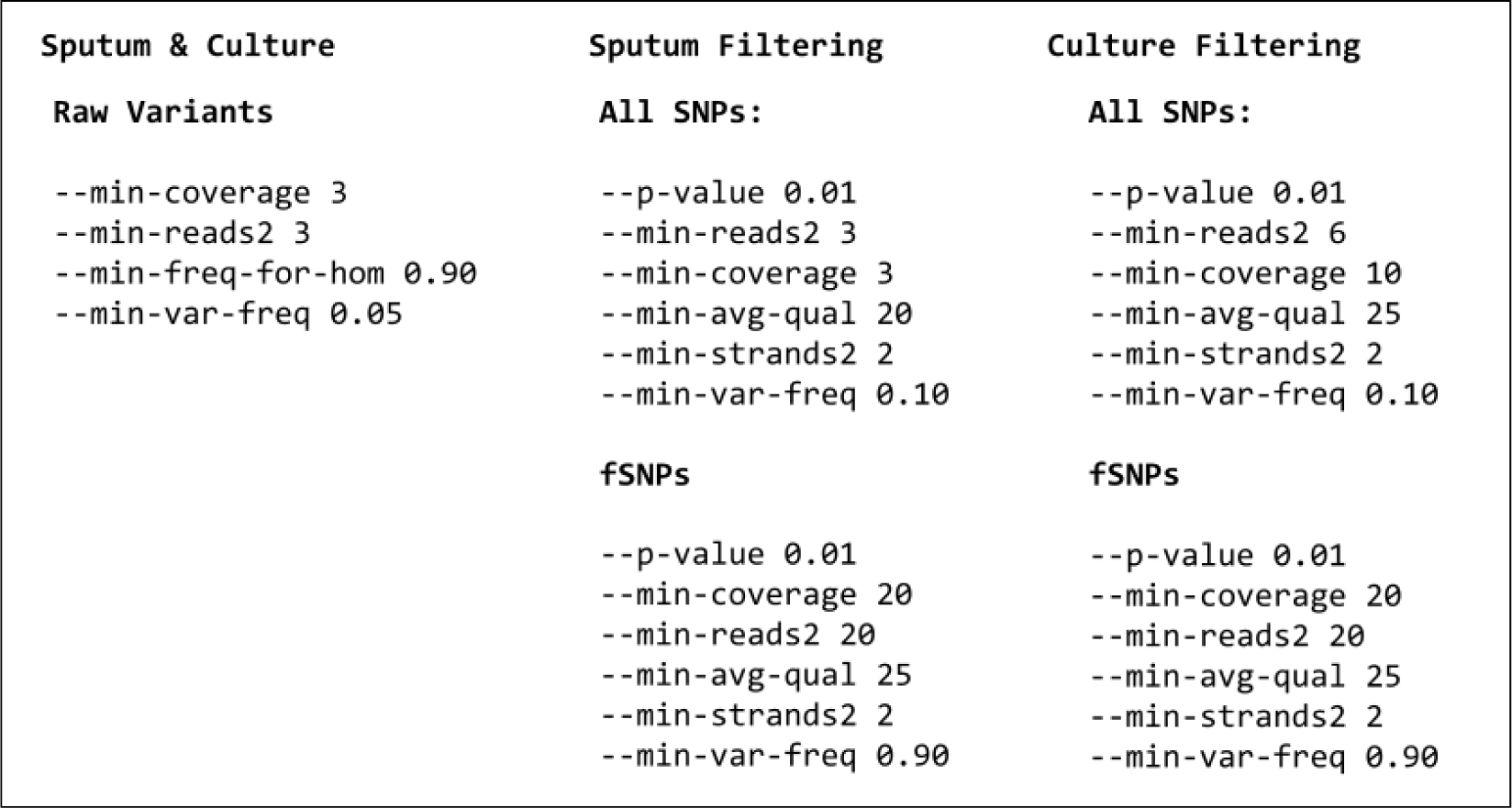
Parameters used for variant calling in VarScan. Fixed SNPs (fSNPs) are the ones called at a frequency above 90%. After obtaining SNP files additional filters were applied as described in the ***Core analysis*** section.

#### Capture technical validation

To investigate if the capture step introduced bias in variant calling, we analysed sputum samples sequenced by direct (dWGS) and enrichment (eWGS) methods and compared to the respective culture (hereafter, trios-analysis). All discrepant positions within trios, but particularly between eWGS and dGWS samples, were manually checked on the reads alignment to the reference genome in Tablet viewer (v1.17.08.17, see **Supplementary material Fig. S2**) to identify the causes of the inconsistency. All discrepant variants were identified in supplementary alignments, which are part of a read that is split and aligned to a different part of the genome than the primary alignment ^18,19^, were identified as the main source of false positive variants. Therefore, we discarded them from bam files running samtools v1.15 (command: samtools view -bh -f0 -F256 -F2048 $BAM_FILE), before variant calling step. Afterwards, we evaluated the impact of discarding supplementary alignments in diversity comparison in all the 61 pairs (sputum-culture) analysed.

After applying all filters described above, a pairwise comparison of variants was conducted within sputum samples and their corresponding cultures. At this point we included a recovery step based on searching for the discrepant SNPs, in the files of the paired sample obtained from a less restrictive variant calling. The reason was that in samples with suboptimal depth the missing SNP may still exist but may have been lost during the filtering steps. Parameters used in VarScan pileup2snp for generating rescued SNP files for comparison analysis were: --min-coverage 2, --min-reads2 1, --min-freq-for-hom 0.9, --min-var-freq 0.01.

The average of rescued SNP shared within sputum and culture pairs was 24 (5-69) representing the 2.5% (0.5-7.9%) of the total variants, highlighting the importance of the step when dealing with samples of heterogeneous coverage.

To verify if rescue was biassing the comparison analysis, we conducted a concordance analysis to corroborate that our rescue approach was not artificially removing true discrepant positions. We analysed 35 sputum-culture pairs with depth of coverage >50x and >95% of the genome covered by applying the same calling pipeline (calling parameters for cultures Methods **Fig. 7**) and skipping the rescue step. Results showed a median of 886 (range: 828-1059) of common variants, 2 (range: 0-6) sputum-exclusive variants and 2 (range: 0-16) culture-exclusive variants. This corresponded to a median Cohen’s Kappa coefficient ^38^ of 0.998 (0.992-1) which represented an almost perfect agreement. In this case, the median Pearson’s correlation of variants frequencies was 0.939 (0.454-0.991).

Finally, variants appearing in both samples were classified as “common”, variants rescued were classified as “common rescued”, sputum-exclusive variants were called “only sputum” and those culture-exclusive were classified as “only culture”.

#### Validation using available datasets

We applied our pipeline described to analyse genetic variability on available sputum-culture paired data sets. We downloaded the sequences under the following project accession numbers from ENA: PRJEB9206 ^12^, PRJNA486713 ^16^ and PRJEB37609 ^10^. Sequences were analysed and only pairs passing the quality filters were compared as explained above (see Bioinformatic Analysis from Methods).

#### Resistance profiling and phylogenetic classification

Resistance profile was obtained by looking for DR-conferring mutations (>10% frequency) listed in the WHO catalogue (version from 03/09/2021) ^39^. Lineage and sublineage classification of the strains was determined by looking for phylogenetic variants reported in bibliography ^20,40^.

Overall, even though the sequencing quality for some sputum was not enough for the comparison analysis, we were able to classify phylogenetically 74/80 (92.5%) sputum samples sequenced and 72/80 (90.0%) at a sublineage level according Coll *et al.* classification ^20^.

Pairwise distance between each sputum-culture pair was obtained using the R package ape ^41^ based on a multiple alignment of fixed SNPs (fSNPs, >90% frequency). In addition, we also obtained the pairwise distance to see whether paired samples clustered together. Neighbour joining trees were constructed using MEGA version X ^42^.

## Ethics

The samples from Mozambique came from a study that was approved by the National Bioethics Committee for Health of Mozambique (CNBS, Ref:369/CNBS/17) and the Internal Bioethics Committee of CISM. Regarding samples coming from Georgia, the ethical approval was obtained from the Institutional Review board (IRB) of the National Center for Tuberculosis and Lung Diseases within the framework of observational clinical study NCT02715271. All methods were performed in accordance with the relevant guidelines and regulations. An informed consent was signed by all participants after providing a verbal explanation and written information about the study. Regarding participants under 18 years of age, the informed consent was obtained from their relatives (parents or guardians). All data were de-identified before the analysis.

## Authors’ Contributions

A. García-Basteiro, B. Saavedra-Cervera and E. Mambuque were responsible for data and samples collection in Mozambique. S. Vashakidze, I. Khurtsilava, Z. Avaliani, A. Rosenthal, A. Gabrielian, M. Shurgaia and N. Shubladze were responsible for data and samples collection in Georgia. C. Mariner-Llicer, M. Torres-Puente, L. Villamayor processed and sequenced the samples. M. Torres-Puente, M. Gabriela and I. Comas conceived and designed the experiments. C. Mariner-Llicer, M. Gabriela and I. Comas were involved in data interpretation. C. Mariner-Llicer, G. Goig, M. Gabriela and I. Comas wrote the paper. All the authors reviewed and approved the manuscript for submission.

## Aknowledgment

This work has been supported by the following: European Research Council (ERC): H2020-ERC-COG/0800; Ministerio Español de Ciencia e Innovación: PID2022-137607OB-I00; Fundació La Caixa: HR21-00415; Stop TB partnership (TB REACH): STBP/TBREACH/GSA/W5-30; and International Science and Technology Center (ISTC): Project #G-2143; National Institute of Allergy and Infectious Diseases (NIH).

We would like to thank Katharine Walter for reviewing the manuscript and providing her feedback.

