## Supplementary material for "Genetic diversity within diagnostic sputum samples is mirrored in the culture of *Mycobacterium tuberculosis*"

#### TITLE

#### CONTENT

##### Supplementary Figures:

**Fig. S1:** Maximum likelihood dendrograms.
**Fig. S2:** View of the reads of one enriched sputum mapped against the reference genome.
**Fig. S3:** Analysis of supplementary alignments of 61 pairs.
**Fig. S4:** Venn Diagrams of SNPs comparisons within sputum- culture pairs.
**Fig. S5:** Comparison of variant frequency in all 61 sputum-culture paired samples.

##### Supplementary Tables

**Table S1:** Information about 61 paired samples sequenced.
**Table S2:** Sublineage classification of sputum-culture paired samples.
**Table S3:** Drug-resistance associated SNPs.

### SUPPLEMENTARY FIGURES

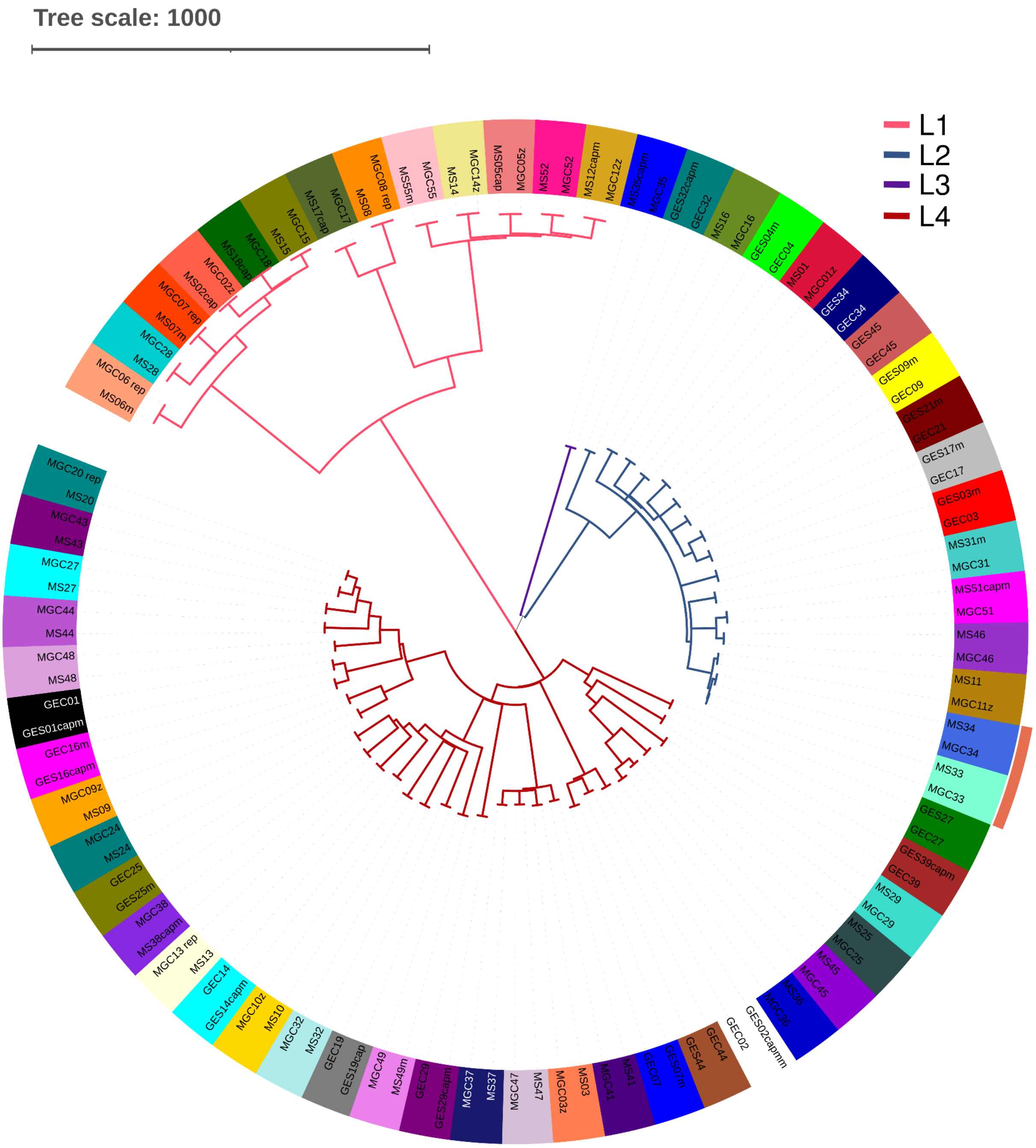


**Fig. S1: Maximum likelihood phylogeny of 61 paired samples.** The colour of the branches represents the lineage of the strains. Paired samples are labelled with the same colour. Transmission clusters are represented with an orange square. Each sputum-culture pair is labelled with a different colour. It was created with Itol (v6.8.1) [(Letunic and Bork 2021)](#).

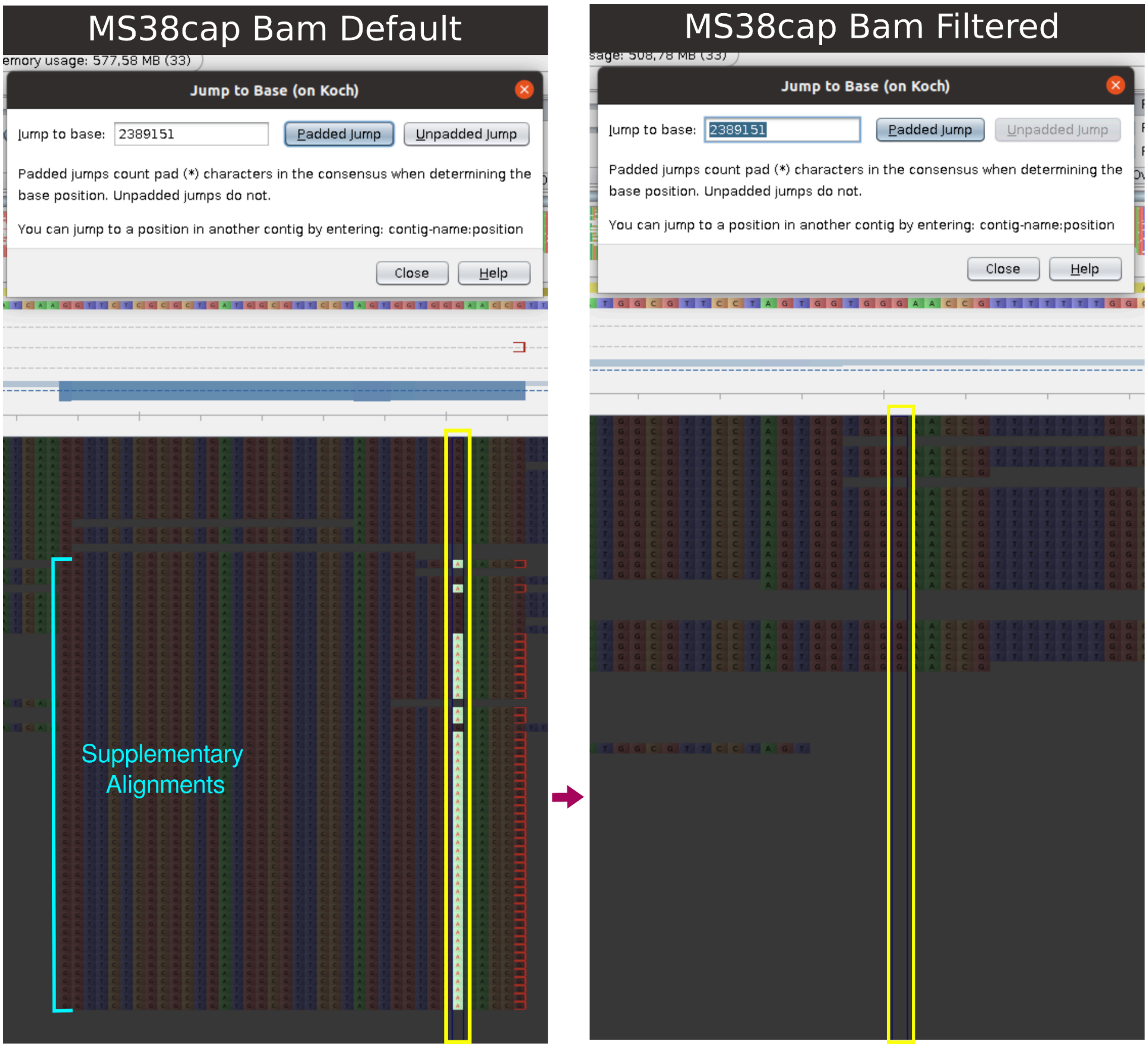


**Fig. S2:** View of the reads of one enriched sputum mapped against the reference genome. The default bam containing the supplementary alignments (labelled in blue) is represented on the left side; and the filtered bam on the right side. The position 2389151 is highlighted in yellow in both alignments showing a false variant in the default bam file at 78.95% frequency that disappeared when discarding the supplementary aligned reads. Images extracted from Tablet viewer.

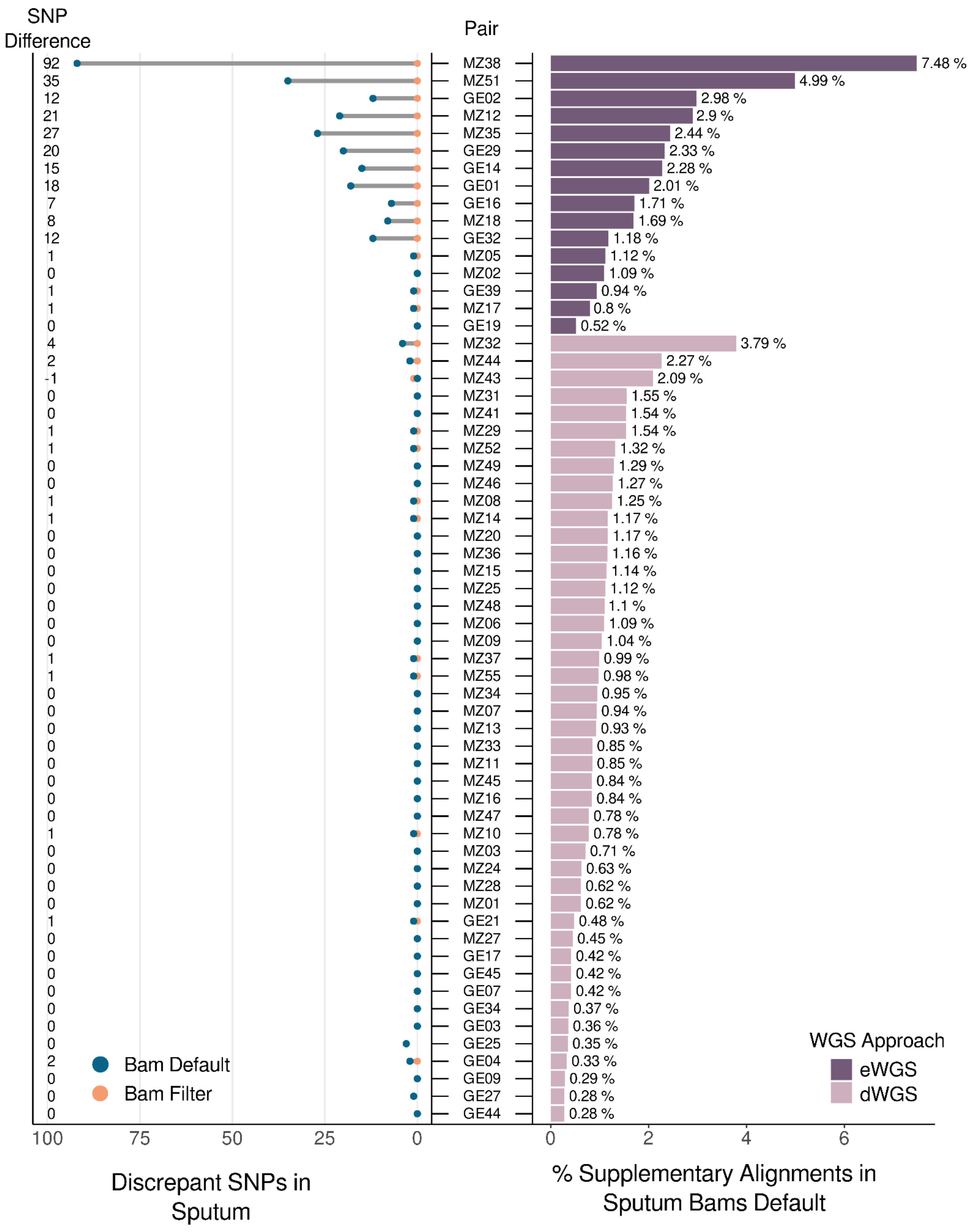


**Fig. S3:** Analysis of supplementary alignments of 61 pairs. The right side represents the percentage of supplementary reads in sputum files. Samples are ordered from the highest to the lowest amount of supplementary reads. Colour represents the sequencing approach. Samples sequenced directly appear in light purple and enriched samples in dark purple. The left side of the plot represents the comparison of the amount of discrepant SNPs only appearing in sputum before and after the bam file filtering. Overlapping points represent a 0 SNP difference. Colours stand for the amount of discrepant variant calls in sputum from bam files before (in blue) and after discarding supplementary reads (in orange).

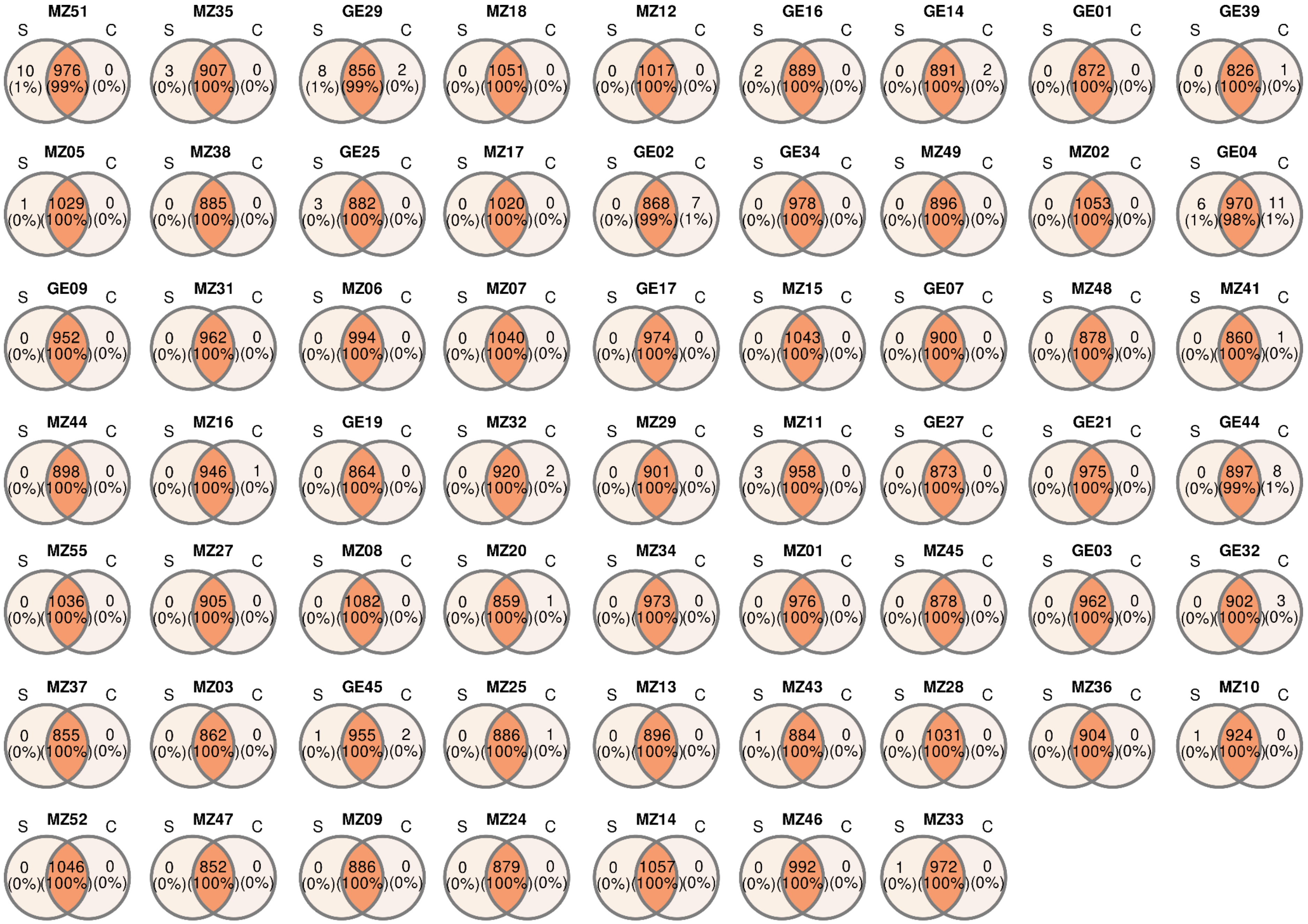


**Fig. S4:** Venn Diagrams of SNPs comparisons within sputum (S) - culture (C) pairs. Supplementary alignments have been discarded for this analysis.

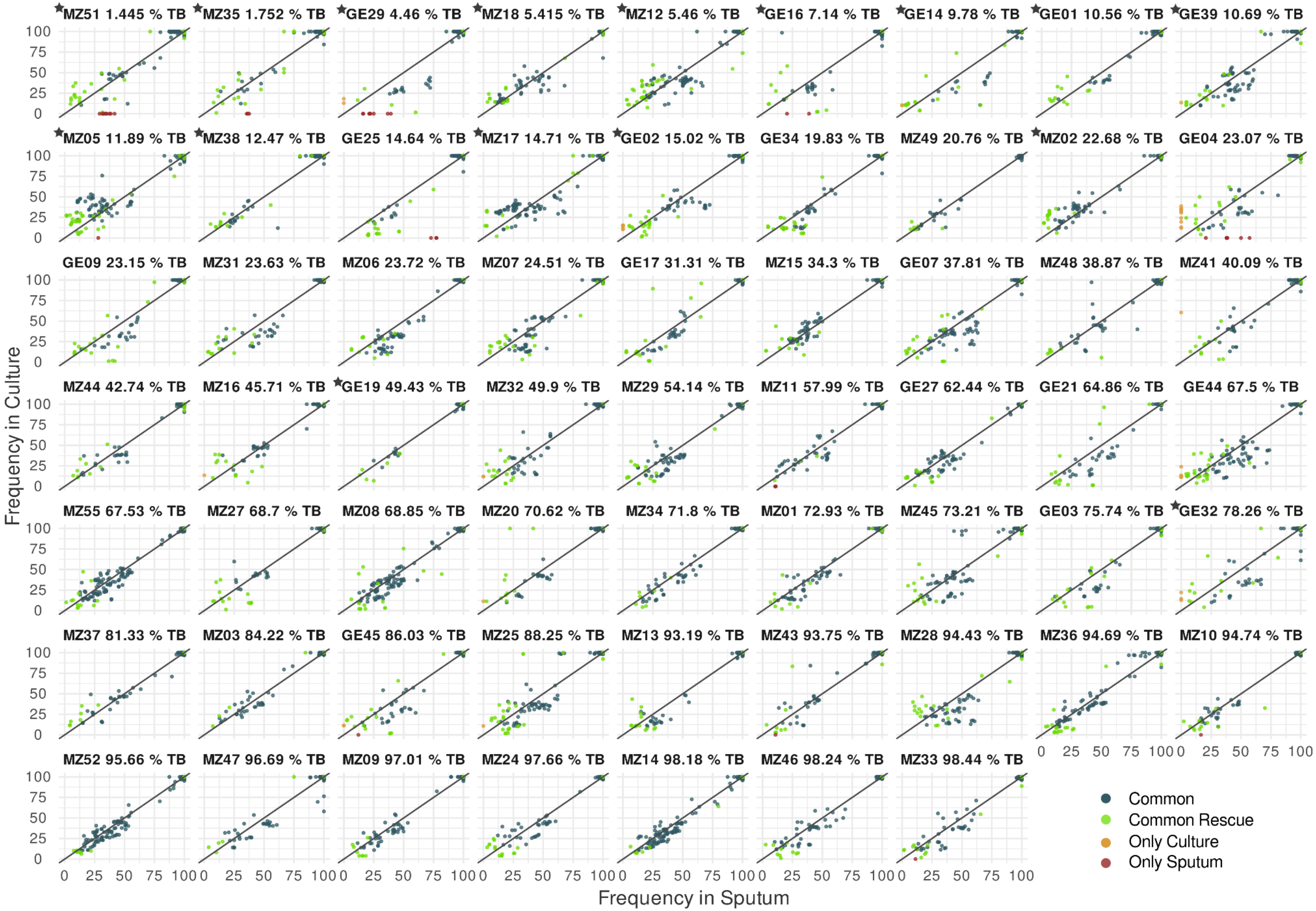


**Fig. S5:** Comparison of variant frequency in all 61 sputum-culture paired samples. Pairs are ordered according to the percentage of MTB in the sputum sample (from the lowest to the highest). Colour of the points represents whether the SNPs are present in both samples (common: blue, common rescued: green), only in the culture (yellow) or only in the sputum (red). The star labelling some pairs indicates that the sputum has been enriched.

### SUPPLEMENTARY TABLES

**Table S1:** information about 61 paired samples sequenced.
**Paired_Samples_61_info_included_Draft**

**Table S2:** Sublineage classification of sputum-culture paired samples.

| Sputum Sample | Phylogeny Sputum | Culture Sample | Phylogeny Culture | Concordance |
|---|---|---|---|---|
| GES01 | lineage4.3.3 | GEC01 | lineage4.3.3 | OK |
| GES02 | lineage4.2.1 | GEC02 | lineage4.2.1 | OK |
| GES03 | lineage2.2.5 | GEC03 | lineage2.2.5 | OK |
| GES04 | lineage2.2.9 | GEC04 | lineage2.2.9 | OK |
| GES07 | lineage4.2.1 | GEC07 | lineage4.2.1 | OK |
| GES09 | lineage2.2.10 | GEC09 | lineage2.2.10 | OK |
| GES14 | lineage4.10 | GEC14 | lineage4.10 | OK |
| GES16 | lineage4.3.3 | GEC16 | lineage4.3.3 | OK |
| GES17 | lineage2.2.10 | GEC17 | lineage2.2.10 | OK |
| GES19 | lineage4.10 | GEC19 | lineage4.10 | OK |
| GES21 | lineage2.2.10 | GEC21 | lineage2.2.10 | OK |
| GES25 | lineage4.10 | GEC25 | lineage4.10 | OK |
| GES27 | lineage4.1.2.1 | GEC27 | lineage4.1.2.1 | OK |
| GES29 | lineage4 | GEC29 | lineage4 | OK |
| GES32 | lineage2.2.2 | GEC32 | lineage2.2.2 | OK |
| GES34 | lineage2.2.10 | GEC34 | lineage2.2.10 | OK |
| GES39 | lineage4.1.1 | GEC39 | lineage4.1.1 | OK |
| GES44 | lineage4.2.1 | GEC44 | lineage4.2.1 | OK |
| GES45 | lineage2.2.10 | GEC45 | lineage2.2.10 | OK |
| MS01 | lineage2.2 | MGC01 | lineage2.2 | OK |
| MS02 | lineage1.1.3 | MGC02 | lineage1.1.3 | OK |
| MS03 | lineage4.4.1.1 | MGC03 | lineage4.4.1.1 | OK |
| MS05 | lineage1.2.2 | MGC05 | lineage1.2.2 | OK |
| MS06 | lineage1.1.3 | MGC06 | lineage1.1.3 | OK |
| MS07 | lineage1.1.3 | MGC07 | lineage1.1.3 | OK |
| MS08 | lineage1.2.2 | MGC08 | lineage1.2.2 | OK |
| MS09 | lineage4.3.2 | MGC09 | lineage4.3.2 | OK |
| MS10 | lineage4.10 | MGC10 | lineage4.10 | OK |
| MS11 | lineage2.2.7 | MGC11 | lineage2.2.7 | OK |
| MS12 | lineage1.2.2 | MGC12 | lineage1.2.2 | OK |
| MS13 | lineage4.10 | MGC13 | lineage4.10 | OK |

| | | | | |
|---|---|---|---|---|
| MS14 | lineage1.2.2 | MGC14 | lineage1.2.2 | OK |
| MS15 | lineage1.1.3 | MGC15 | lineage1.1.3 | OK |
| MS16 | lineage2.2 | MGC16 | lineage2.2 | OK |
| MS17 | lineage1.2.2 | MGC17 | lineage1.2.2 | OK |
| MS18 | lineage1.1.3 | MGC18 | lineage1.1.3 | OK |
| MS20 | lineage4.3.4.2.1 | MGC20 | lineage4.3.4.2.1 | OK |
| MS24 | lineage4.3.2.1 | MGC24 | lineage4.3.2.1 | OK |
| MS25 | lineage4.1.1.3 | MGC25 | lineage4.1.1.3 | OK |
| MS27 | lineage4.3.4.2.1 | MGC27 | lineage4.3.4.2.1 | OK |
| MS28 | lineage1.1.3 | MGC28 | lineage1.1.3 | OK |
| MS29 | lineage4.1.1.2 | MGC29 | lineage4.1.1.2 | OK |
| MS31 | lineage2.2.7 | MGC31 | lineage2.2.7 | OK |
| MS32 | lineage4.10 | MGC32 | lineage4.10 | OK |
| MS33 | lineage2.2.7 | MGC33 | lineage2.2.7 | OK |
| MS34 | lineage2.2.7 | MGC34 | lineage2.2.7 | OK |
| MS35 | lineage3 | MGC35 | lineage3 | OK |
| MS36 | lineage4.1.1.3 | MGC36 | lineage4.1.1.3 | OK |
| MS37 | lineage4.4.1.1 | MGC37 | lineage4.4.1.1 | OK |
| MS38 | lineage4.10 | MGC38 | lineage4.10 | OK |
| MS41 | lineage4.4.1.1 | MGC41 | lineage4.4.1.1 | OK |
| MS43 | lineage4.3.4.2.1 | MGC43 | lineage4.3.4.2.1 | OK |
| MS44 | lineage4.3.4.2 | MGC44 | lineage4.3.4.2 | OK |
| MS45 | lineage4.1.1.3 | MGC45 | lineage4.1.1.3 | OK |
| MS46 | lineage2.2.7 | MGC46 | lineage2.2.7 | OK |
| MS47 | lineage4.4.1.1 | MGC47 | lineage4.4.1.1 | OK |
| MS48 | lineage4.3.4.1 | MGC48 | lineage4.3.4.1 | OK |
| MS49 | lineage4.10 | MGC49 | lineage4.10 | OK |
| MS51 | lineage2.2.7 | MGC51 | lineage2.2.7 | OK |
| MS52 | lineage1.2.2 | MGC52 | lineage1.2.2 | OK |
| MS55 | lineage1.2.2 | MGC55 | lineage1.2.2 | OK |

**Table S3:** Drug-resistance associated SNPs.

| Pair | Gene | Rv Gene | Genomic Position | WT codon | Mut Codon | aa Change | Antibiotic | WHO Classification | Frequency Sputum (%) | Frequency Culture (%) |
|---|---|---|---|---|---|---|---|---|---|---|
| GE01 | katG | Rv1908c | 2155168 | AGC | ACC | S315T | INH | Assoc w R | 100 | 100 |
| GE02 | fabG1_inhA | IG_Rv1482c_Rv1483 | 1673425 | C | T | c-777t | INH, ETH | Assoc w R | 100 | 100 |
| GE04* | rpoB | Rv0667 | 761155 | TCG | TTG | S450L | RIF | Assoc w R | 100 | 100 |
| | katG | Rv1908c | 2155168 | AGC | ACC | S315T | INH | Assoc w R | 100 | 100 |
| | eis_IG | IG_Rv2416c_Rv2417c | 2715369 | C | A | g-37t | KAN | Assoc w R | 100 | 100 |
| | embB | Rv3795 | 4248003 | CAG | CGG | Q497R | EMB | Assoc w R | 100 | 100 |
| | ethA | Rv3854c | 4326707 | TGG | TAG | W256! | ETH | Assoc w R - Interim | 100 | 100 |
| GE29 | katG | Rv1908c | 2155168 | AGC | ACC | S315T | INH | Assoc w R | 100 | 100 |
| GE32** | gyrA | Rv0006 | 7581 | GAC | TAC | D94Y | MXF, LEV | Assoc w R | 100 | 100 |
| | rpoB | Rv0667 | 761155 | TCG | TTG | S450L | RIF | Assoc w R | 100 | 100 |
| | katG | Rv1908c | 2155168 | AGC | ACC | S315T | INH | Assoc w R | 94.74 | 100 |
| | embB | Rv3795 | 4248003 | CAG | CGG | Q497R | EMB | Assoc w R | 100 | 100 |
| GE39 | katG | Rv1908c | 2155168 | AGC | ACC | S315T | INH | Assoc w R | 100 | 100 |
| MZ07 | katG | Rv1908c | 2155168 | AGC | ACC | S315T | INH | Assoc w R | 100 | 100 |
| MZ09 | katG | Rv1908c | 2155168 | AGC | ACC | S315T | INH | Assoc w R | 100 | 100 |
| MZ12 | fabG1_inhA | IG_Rv1482c_Rv1483 | 1673425 | C | T | c-777t | INH, ETH | Assoc w R | 100 | 100 |
| MZ14* | fabG1_inhA | IG_Rv1482c_Rv1483 | 1673425 | C | T | c-777t | INH, ETH | Assoc w R | 100 | 100 |
| | rpoB | Rv0667 | 761155 | TCG | TTG | S450L | RIF | Assoc w R | 100 | 100 |
| | katG | Rv1908c | 2155168 | AGC | ACC | S315T | INH | Assoc w R | 100 | 100 |
| | pncA | Rv2043c | 2288826 | GTG | GGG | V139G | PZA | Assoc w R | 100 | 100 |
| | embB | Rv3795 | 4247431 | ATG | ATA | M306I | EMB | Assoc w R | 100 | 98.59 |

**Abbreviations:** IG - Intergenic Region, INH - isoniazid, ETH - ethionamide, SM - streptomycin, EMB - ethambutol, ETH - ethionamide, KAN - kanamycin, RMP - rifampicin, FQ - fluoroquinolones, PZA - pyrazinamide, ! - Stop codon
